# 3D-bioprinted, phototunable hydrogel models for studying adventitial fibroblast activation in pulmonary arterial hypertension

**DOI:** 10.1101/2022.11.11.516188

**Authors:** Duncan Davis-Hall, Emily Thomas, Brisa Peña, Chelsea M. Magin

## Abstract

Pulmonary arterial hypertension (PAH) is a progressive disease of the lung vasculature, characterized by elevated pulmonary blood pressure, remodeling of the pulmonary arteries, and ultimately right ventricular failure. Therapeutic interventions for PAH are limited in part by the lack of *in vitro* screening platforms that accurately reproduce dynamic arterial wall mechanical properties. Here we present a 3D-bioprinted model of the pulmonary arterial adventitia comprised of a phototunable poly(ethylene glycol) alpha methacrylate (PEG-αMA)-based hydrogel and primary human pulmonary artery adventitia fibroblasts (HPAAFs). This unique biomaterial emulates PAH pathogenesis *in vitro* through a two-step polymerization reaction. First, PEG-αMA macromer was crosslinked off-stoichiometry by 3D bioprinting an acidic bioink solution into a basic gelatin support bath initiating a base-catalyzed thiol-ene reaction with synthetic and biodegradable crosslinkers. Then, matrix stiffening was induced by photoinitiated homopolymerization of unreacted αMA end groups. A design of experiments approach produced a hydrogel platform that exhibited an initial elastic modulus (E) within the range of healthy pulmonary arterial tissue (E = 4.7 ± 0.09 kPa) that was stiffened to the pathologic range of hypertensive tissue (E = 12.8 ± 0.47 kPa) and supported cellular proliferation over time. A higher percentage of HPAAFs cultured in stiffened hydrogels expressed the fibrotic marker alpha-smooth muscle actin than cells in soft hydrogels (88 ± 2% versus 65 ± 4%). Likewise, a greater percentage of HPAAFs were positive for the proliferation marker 5-ethynyl-2’-deoxyuridine (EdU) in stiffened models (66 ± 6%) compared to soft (39 ± 6%). These results demonstrate that 3D-bioprinted, phototunable models of pulmonary artery adventitia are a tool that enable investigation of fibrotic pathogenesis *in vitro*.

## Introduction

Pulmonary arterial hypertension (PAH) is a chronic disease characterized by abnormal and progressive cellular proliferation, phenotype dysregulation, and extracellular matrix (ECM) remodeling, which leads to increased blood pressure in the arteries of the lungs and eventually heart failure.^[1]^ The pulmonary arteries consist of three layers. From inner to outermost these layers are the tunica intima, the tunica media, and the tunica adventitia.^[2]^ It has long been understood that the endothelial cells that line the tunica intima and the smooth muscle cells that reside in the tunica media play a major role in PAH pathology.^[3, 4]^ The role of the tunica adventitia in PAH pathophysiology is just beginning to emerge. In this study, we designed and evaluated a 3D-bioprinted model of pulmonary arterial adventitia that uses a two-step polymerization process to increase matrix modulus with precise control in time and space around primary human pulmonary artery adventitia fibroblasts (HPAAFs).

Recent evidence indicates a causative relationship between ECM remodeling in the tunica adventitia and activation of the fibroblasts that reside within to a myofibroblast phenotype.^[5-7]^ Adventitial ECM alterations and the subsequent increase in tissue stiffness – from an elastic modulus (E) of 1-5 kPa in healthy tissue to more than 10 kPa in diseased tissue^[8]^ – are suggested to be an early and potent driving force for the continuous alteration of fibroblast phenotype and function, as well as a contributor to the physical alteration of pulmonary circulation.^[9]^ Activated myofibroblasts express the intracellular contractile protein alpha-smooth muscle actin (αSMA) and exhibit increased collagen production to promote ECM remodeling and wound healing. These cells typically undergo apoptosis once healing is complete in healthy tissues to maintain homeostasis.^[10, 11]^ However, chronically activated myofibroblasts that resist apoptosis and continue to remodel the surrounding ECM have been identified within hypertensive pulmonary arteries.^[12-14]^ These cells continually deposit new collagen, initiate collagen crosslinking, and contribute to elastic laminae degradation within the vessel wall.^[15]^ Based on experiments performed in two-dimensional (2D) *in vitro* models of fibrosis, persistent fibroblast activation could be tied to a self-perpetuating cycle where fibrotic ECM locally induces fibrotic activity in nearby fibroblasts leading to chronic and progressive fibrosis.^[16]^ Therapeutic approaches to reverse this perpetual fibrotic activation are not yet clinically available.^[14]^

Leveraging biomaterial design and advanced biofabrication techniques to build tools that improve our understanding of how changes in elastic modulus within a three-dimensional (3D) microenvironment influence fibroblast activation may illuminate possible therapeutic targets for PAH. There is a growing class of dynamic hydrogel biomaterials that exploits sequential polymerization reactions to allow user-controlled manipulation of microenvironmental mechanical properties. These systems have been engineered to control and study 3D cell-matrix interactions in real time.^[17-21]^ Günay et al. used dynamically stiffening anthracene-functionalized poly(ethylene glycol) (PEG) hydrogels to investigate cardiac fibroblast mechanobiology. This study measured fibroblast expression of nuclear factor of activated T cells (NFAT), which localizes in cell nuclei following intracellular calcium signaling and initiates transcription of fibrotic genes. Static hydrogels with elastic moduli of 10 kPa and 50 kPa were used as cellular substrates, along with dynamic hydrogels that could stiffen from 10 to 50 kPa. The results of this study indicated that dynamic substrate stiffening increased NFAT nuclear localization in cardiac fibroblasts, indicating fibrotic activation.^[17]^ Another study measured the effects of a stiffening 3D PEG-based hydrogel on the phenotype of valvular interstitial cells (VICs), another cardiac fibroblast. VICs cultured in 3D hydrogels that stiffened from 0.24 to 13 kPa expressed only 23% as much αSMA, an indicator of myofibroblast activation, after stiffening.^[18]^ These results differed from the observed correlation between substrate stiffness and fibrotic activation seen in 2D, suggesting that culture dimensionality plays an important role in cell behavior.^[22]^ Other groups have recently used 3D bioprinting to fabricate models of lung structures.^[23-25]^ Polyvinylpyrrolidone bioink laden with human lung epithelial cells, endothelial cells, and fibroblasts was used to create a multicellular alveolar construct. This alveolar model maintained cell viability and demonstrated similar proliferation to non-printed cells.^[23]^ Another group 3D bioprinted pulmonary artery anastomosis models from gelatin methacrylate hydrogels seeded with human umbilical vein endothelial cells – cultured under physiologic flow conditions, these constructs sustained cell viability.^[25]^ These studies demonstrate that pulmonary 3D bioprinting techniques have seen major advancement.

To build a tool for investigating the effects of dynamic stiffening on fibrotic activation in 3D, we present a biomaterial and biofabrication approach that recapitulate the geometry of the pulmonary arterial adventitia and allows the user to control mechanical properties in real time. A phototunable PEG-alpha methacrylate (PEG-αMA) hydrogel was fine-tuned for 3D bioprinting these models using the freeform reversible embedding of soft hydrogels (FRESH) technique.^[26]^ Results showed that it provided a hydrolytically stable microenvironment with dynamically tunable mechanical properties. Hydrogels were initially formulated to reproduce the elastic modulus of healthy pulmonary arterial adventitia and stiffened to mimic hypertensive tissue. HPAAFs cultured in stiffened hydrogel bioprints showed a higher percentage of αSMA- and 5-ethynyl-2’-deoxyuridine (EdU)-positive cells, indicating activation, compared to HPAAFs grown in soft 3D hydrogels. The model presented here is a versatile new tool for studying fibroblast activation in the pulmonary arterial adventitia that could provide critical mechanistic insights into PAH pathology and treatment.

## Materials and Methods

### Synthesis of PEG-αMA

PEG-αMA was synthesized as previously described.^[27]^ Briefly, poly(ethylene glycol)-hydroxyl (PEG-OH; 8-arm, 10 kg mol^−1^; JenKem Technology, Plano, TX) was dissolved in anhydrous tetrahydrofuran (THF; Sigma-Aldrich, St. Louis, MO) in a round-bottom flask and purged with argon. Sodium hydride (NaH; Sigma-Aldrich, St. Louis, MO) was dissolved in THF and injected through a septum into the reaction vessel at 3× molar excess to PEG-OH groups.^[28]^ Ethyl 2-(bromomethyl)acrylate (EBrMA; synthesized as previously described)^[27]^ was added drop-wise using an addition funnel at a 6× molar ratio to PEG-OH groups, and the reaction was stirred at room temperature for 72 hours protected from light. The mixture was neutralized with 1N acetic acid until gas evolution ceased and vacuum filtered through Celite 545. The solution was concentrated by rotary evaporation at 60 °C, precipitated dropwise into ice-cold diethyl ether (Thermo Fisher Scientific, Waltham, MA), and washed three times through centrifugation in diethyl ether. The solid precipitate was then dried under vacuum overnight at room temperature. The product was purified using dialysis (1 kg mol^−1^ MWCO, Thermo Fisher Scientific, Waltham, MA) for four days, and then flash frozen in liquid nitrogen and lyophilized to give the final product.^[27]^ The functionalization of the product was verified by ^1^H NMR.

### Synthesis of PEGMA

PEGMA was synthesized as previously described and characterized using ^1^H NMR.^[29, 30]^

### Rheological Characterization

The influence of 10 kg mol^−1^ PEG-αMA weight percent (wt%) and the ratio of crosslinking agents, dithiothreitol (DTT; Sigma-Aldrich, St. Louis, MO) to a matrix metalloproteinase 2 (MMP2)-degradable peptide sequence (KCGGPQGIWGQGCK; GL Biochem, Boston, MA) on hydrogel elastic modulus was investigated with a design of experiments (DOE) approach. Two crosslinkers, including the relatively low molecular weight DTT, were necessary to achieve the desired increase in hydrogel mechanical properties while enabling fibroblasts to remodel the microenvironment through MMP2 enzymatic degradation to replicate spreading and activation *in vivo*.^[31]^ Adventitial fibroblasts produce MMP2 *in vivo* during healthy tissue regulation and diseased states of vascular remodeling.^[32]^ The peptide sequence chosen for this platform was optimized for degradation by MMP2.^[33]^ The DOE statistical approach identified a multifactorial experiment that systematically determined the individual and combinatorial effects of each variable on elastic modulus and cell proliferation. Hydrogels with varying PEG-αMA wt% and DTT:MMP2-degradable crosslinker ratios (**Table 1**) were prepared with pH 8.0 N-2-hydroxyethylpiperaine-N-2-ethane sulfonic acid (HEPES; Thermo Fisher Scientific, Waltham, MA) as a solvent to allow base-catalyzed polymerization. Prepared hydrogels were permitted to swell to equilibrium in a 2.2 mM lithium phenyl-2,4,6-trimethylbenzoylphosphinate (LAP; synthesized as previously described)^[34, 35]^ solution in phosphate buffered saline (PBS; Life Technologies, Carlsbad, CA) to introduce photoinitiator for the stiffening reaction.

**Table 1.** Hydrogel formulations for rheological characterization.

| **Platform** | **DTT:MMP2-degradable crosslinker** | **PEG-αMA wt%** |
| --- | --- | --- |
| **1** | 0.9:0.1 | 15 |
| **2** | 0.9:0.1 | 17.5 |
| **3** | 0.7:0.3 | 12.5 |
| **4** | 0.7:0.3 | 17.5 |
| **5** | 0.5:0.5 | 12.5 |
| **6** | 0.5:0.5 | 15 |
| 7 | 0.5:0.5 | 17.5 |

Storage modulus (G’) of 8-mm diameter circular punched hydrogels was measured with parallel plate rheology using oscillatory shear at 1% strain through a dynamic angular frequency range of 0.1 to 100 rad s^−1^ on a Discovery Hybrid Rheometer 2 (DHR-2; TA Instruments, New Castle, DE). Hydrogel samples were placed on the rheometer’s Peltier plate set to 37 °C. An 8-mm parallel plate geometry was lowered onto disc samples until an axial force of 0.03 N was achieved – this gap distance was considered initial contact, or 0% compression. Gap distance was lowered in successive increments and elastic modulus was measured until G’ plateaued for each hydrogel formulation. The plateau gap distance was expressed as a percent compression and was used for successive measurements of that hydrogel formulation.^[36]^ Hydrogel stiffening was achieved by photopolymerization of all samples initiated by exposure to 10 mW cm^−2^ 365-nm light (OmniCure Series 2000, Lumen Dynamics, Monroe, LA) for five minutes. Post-stiffening changes to G’ were measured. Elastic modulus was calculated from storage modulus, E = 2G’(1 + ν), in the linear viscoelastic region assuming Poisson’s ratio of an incompressible material, 0.5.^[27, 37]^

Real-time rheology was conducted to measure the elastic moduli of PEG-αMA hydrogels through both stages of polymerization: the initial base-catalyzed polymerization and the subsequent photoinitiated homopolymerization. Hydrogel precursor solution for real-time rheology contained 17.5 wt% PEG-αMA polymer solution with a 0.375 molar ratio relative to PEG-αMA backbone of DTT and MMP2-degradable crosslinker (70:30 ratio of crosslinkers, respectively) dissolved in pH 8.0 HEPES. LAP (2.2 mM in HEPES) was included in the hydrogel precursor solution to allow photoinitiated stiffening. Immediately after combining hydrogel components, 50 µL of liquid hydrogel precursor solution was deposited on the rheometer Peltier plate set to 37 °C and the 8-mm parallel plate geometry was lowered until it contacted the liquid sample. Storage modulus was measured at a constant 10 rad s^−1^ at 1% strain for 12 minutes. After five minutes of measurements, samples were stiffened by exposure to 10 mW cm^−2^ 365-nm light for five minutes. Elastic modulus was calculated from G’ using the equation above.

### Hydrogel Formation for 2D Metabolic Activity Studies

Hydrogel precursor solutions prepared with 2 mM CGRGDS (RGD; GL Biochem, Boston, MA), 8-arm, 10 kg mol^−1^ PEG-αMA, and the DTT:MMP2-degradable crosslinker ratios described in **Table 2** with a 0.375 ratio of crosslinker to PEG-αMA backbone in sterile pH 8.0 HEPES. Glass coverslips were coated with (3-mercaptopropyl) trimethoxysilane (Acros Organics, Fair Lawn, NJ) following a liquid deposition silanation technique.^[38]^ Hydrogel precursor (90 µL) was placed between a Sigmacote (Sigma-Aldrich, St. Louis, MO)-treated glass microscope slide and a silanated glass coverslip. Initial polymerization occurred spontaneously using pH 8.0 HEPES as solvent.

**Table 2.** Hydrogel formulations for assessment of 2D cellular metabolic activity.

| Platform | DTT:MMP2-degradable crosslinker | PEG-αMA wt% |
| --- | --- | --- |
| 1 | 0.9:0.1 | 15 |
| 2 | 0.7:0.3 | 17.5 |
| 3 | 0.5:0.5 | 15 |
| 4 | 0.5:0.5 | 17.5 |

Polymerized hydrogels were swollen in media made from Dulbecco’s Modified Eagle Medium (DMEM)/F12 1:1 supplemented with 10% fetal bovine serum (FBS), 100 U mL^−1^ penicillin, 100 µg mL^−1^ streptomycin, and 2.5 µg mL^−1^ amphotericin B (Gibco), with 2.2 mM LAP overnight at 37 °C. Stiffening was initiated by exposure to 10 mW cm^−2^ 365-nm light for five minutes.

Primary human lung fibroblast (HLF; passage 3; ATCC PCS-201-013, Manassas, VA) metabolic activity was assessed using PrestoBlue Cell Viability Reagent (Thermo Fisher Scientific, Waltham, MA) on days 1, 3, 5, and 7 per manufacturer’s instructions.

### Cell Culture Methods

HLFs were cultured in growth media made from DMEM/F12 1:1 (Gibco, Waltham, MA) supplemented with 10% FBS, 50 U mL^−1^ penicillin, 50 × 10^−6^ M mL^−1^ streptomycin, and 0.5 µg mL^−1^ amphotericin B (Sigma-Aldrich, St. Louis, MO) for metabolic activity experiments.^[37]^ Human pulmonary artery adventitia fibroblasts (HPAAFs; Accegen, Fairfield, NJ) from a 2-year-old male patient were cultured in complete HPAAF culture media according to manufacturer instructions. Complete media consisted of basal media supplemented with 10% FBS, 2% fibroblast growth serum (FGS), 50 U mL^−1^ penicillin, 50 × 10^−6^ M mL^−1^ streptomycin (Accegen), and 0.5 µg mL^−1^ amphotericin B (Sigma-Aldrich).

### Hydrolytic Stability

PEG-αMA hydrogels were fabricated at 17.5 wt% with a 0.375 molar ratio of 70:30 DTT:MMP2-degradable crosslinker as described above to measure hydrolytic stability compared to widely used PEG-methacrylate (PEG-MA; 8-arm, 10 kg mol^−1^) hydrogels. PEG-MA hydrogels were fabricated at 19 wt% and an equivalent crosslinker composition to match the initial elastic modulus of PEG-αMA hydrogels. All hydrogel samples were swollen in 2.2 mM LAP in PBS overnight and then stiffened with 10 mW cm^−2^ 365-nm light for five minutes. Samples were then soaked in PBS at 37 °C and elastic modulus was measured at days 1, 10, 20, 40, and 60.

### Patterning of Hydrogel Elastic Modulus

PEG-αMA hydrogels were fabricated at 17.5 wt% with a 0.375 molar ratio of 70:30 DTT:MMP2-degradable as described above and swollen overnight in PBS with 2.2 mM LAP and 10 µM methacryloxyethyl thiocarbamoyl rhodamine B (Polysciences Inc., Warrington, PA). Swollen hydrogels were exposed to 10 mW cm^−2^ 365-nm light for five minutes through a chrome-on-quartz photomask with spatially patterned opaque and transparent regions. The photomask had a pattern of 100-µm wide transparent stripes spaced by 100-µm wide opaque regions. Stiffened hydrogels were soaked in PBS at 4 °C for two days to remove excess rhodamine dye and then imaged with a TRITC filter to reveal stiffened regions.

### Atomic Force Microscopy

Hydrogel precursor solution (17.5 wt% PEG-αMA with a 0.375 molar ratio of 70:30 DTT:MMP2-degradable crosslinker) was deposited in 40 µL drops between a silanated glass coverslip (25 × 75 mm #1 thickness; Bellco Glass) and a hydrophobic glass microscope slide (25 × 75 mm; Fisher Scientific) separated by a 0.5-mm tall silicone gasket to create thin, flat hydrogel discs for atomic force microscopy (AFM) measurement. Polymerized hydrogel samples were photopatterned with 100-µm wide stiffened stripes as described above, but without the addition of methacryloxyethyl thiocarbamoyl rhodamine B. Photopatterned hydrogel samples were soaked overnight in PBS and then elastic moduli of soft and stiffened stripes were measured with AFM (JPK NanoWizard 4a BioScience). Force Spectroscopy mode was used to determine the nanomechanical properties of the polymers. Silicon nitride AFM probes (MLCT; Bruker, Billerica, MA) were used with a pyramidal Si_3_N_4_ 35° curvature radius tip. The spring constants of all cantilevers were systematically measured (Bruker: 0.1-0.2 N m^−1^) using the thermal tune method. The maximal force applied to the polymers was limited to 1.5 nN, with 5 µm of Z piezo. All physical cues regarding AFM analysis and sample preparation were kept constant across all samples. The sections across the polymers were monitored and their morphological details were observed using an optical light microscope. Polymer upper, middle, and bottom sections denoted three different photopatterned stripes. An average of 15 measurements per polymer section from two polymers were analyzed. The Hertz model was used to obtain the elasticity of the polymers using JPKSPM Data Processing software (JPK).

### 3D Bioprinting

All 3D bioprinting was performed using the FRESH technique and a custom-designed 3D bioprinter. Briefly, a LulzBot Mini 2 (Aleph Objects, Inc.) was modified with a custom-built syringe pump extruder that replaced the stock thermoplastic extruder to convert this hardware into a bioprinter.^[39]^ 3D computer-aided design (CAD) models were created in Fusion 360, then sliced into G-code using Slic3r software. This G-code was uploaded to Pronterface, which interfaced with the modified Mini Lulzbot 2 3D printer to provide hardware control. Pronterface software sent 3D bioprinting instructions to the bioprinter to create cylinders with 4-mm height and 4-mm inner diameter and variable wall thicknesses (**Table 3**). To improve shear-thinning and reduce cell settling, 2.5 wt% 400 kg mol^−1^ poly(ethylene oxide) (PEO) was added to the final hydrogel formulation (supplementary information). This bioink and HPAAFs were loaded into a glass syringe with a 150-μm (inside diameter) stainless steel needle tip mounted in the new extruder. This bioink solution was kept at pH 6.2 to prevent base-catalyzed polymerization in the extruder during bioprinting. Wells of a 24-well plate were filled with LifeSupport gelatin microparticle slurry (Advanced BioMatrix) and mounted on the 3D bioprinter build plate. The LifeSupport slurry was maintained at pH 9.0 to initiate base-catalyzed polymerization of bioink after deposition in the support bath. The tip of the syringe needle was positioned at the center of the gelatin support bath in x and y and at least 1 mm above the bottom of the bath in the z-direction. The bioink was extruded through a thin needle by a syringe pump into a gelatin microparticle support bath where it polymerized. Sixty minutes after bioprinting, the constructs were heated to 37 ºC in an incubator overnight to melt and remove the gelatin support bath (**Fig S1A**). Melted gelatin support was exchanged for complete media the day after bioprinting. 3D-bioprinted constructs were cultured in complete human pulmonary artery adventitia fibroblast medium (AcceGen) supplemented with 2% FBS, 50 U mL^−1^ penicillin, 50 × 10^−6^ M mL^−1^ streptomycin, and 0.5 μg mL^−1^ amphotericin B (Sigma) for metabolic activity experiments.^[37]^

**Table 3.** Experimental conditions for evaluation of 3D cell viability.

| Platform | Wall Thickness ($\mu\text{m}$)<sup>[40, 41]</sup> | Cell Density (cells $\text{mL}^{-1}$)<sup>[41, 42]</sup> |
| --- | --- | --- |
| 1 | 300 | $4 \times 10^5$ |
| 2 | 300 | $4 \times 10^6$ |
| **3** | 500 | $4 \times 10^5$ |
| **4** | 500 | $4 \times 10^6$ |

### 3D Cell Viability

3D-bioprinted constructs were fabricated as described above with different wall thicknesses and HPAAF densities (Table 3, n = 3 for each condition). LifeSupport gelatin microparticle slurry was exchanged for complete media the day after bioprinting (day 1) and media was changed every three days after until each time point for viability assessment. Constructs were stained with a Live/Dead Double Staining Kit (EMD Millipore, Burlington, MA) according to manufacturer instructions at days 1, 3, 7, 14, and 21 following bioprinting. Briefly, constructs were rinsed in sterile PBS and then incubated with Live/Dead solution on a rocker at 37 ºC for 40 minutes. The Live/Dead solution consisted of a 1:1,000 dilution of calcein AM for live cells and propidium iodide for dead cells in staining buffer. Constructs were then transferred to sterile PBS and imaged immediately on a Keyence BZ-X810 all-in-one fluorescence microscope (Keyence, Osaka, Japan). Viability was expressed as average percentage of live cells from three different 100-µm z-stacks each on three constructs per condition per time point.

### Assessment of Activation by Immunofluorescent Staining

Soft arterial adventitia models with 300-µm wall thickness and 4 × 10^6^ HPAAFs mL^−1^ were FRESH 3D-bioprinted as described above. LifeSupport microparticle slurry was exchanged for activation media (complete HPAAF media as described above, containing 1% FBS) the day after bioprinting (day 1) and media was changed again on day 4. Activation media was changed again on day 6 and supplemented with 2.2 mM sterile LAP photoinitiator. On day 7, half of the soft samples (n = 3) were removed from media and rinsed with sterile PBS. These samples were fixed with 4% paraformaldehyde (PFA; Electron Microscopy Sciences, Hatfield, PA) in sterile PBS for 30 minutes on an orbital rocker at 37 ºC. Samples were then quenched with 100 mM glycine (Sigma) in PBS for 15 minutes on a rocker at room temperature and soaked in optimal cutting temperature compound (OCT; Fisher Scientific) at 4 ºC overnight. Also on day 7, the remaining half of the soft samples (n = 3) were stiffened with 10 mW cm^−2^ 365-nm light for five minutes and placed in fresh activation media. On day 8, soft OCT-soaked samples were flash-frozen in cryomolds containing additional OCT in a plastic vessel placed inside another vessel containing 2-methylbutane (Fisher Scientific) placed inside a Styrofoam box containing liquid nitrogen. Stiffened samples were fixed, quenched, and soaked in OCT as described above on day 9, then frozen as described above on day 10. All OCT-embedded samples were stored at -80 ºC until cryosectioning.

Frozen samples were cryosectioned at -22 ºC. Slices (10-µm thick) were taken from each sample and attached to positively charged glass microscope slides (Globe Scientific, Mahwah, NJ), with three glass microscope slides covered with 10-12 cryosections per 3D hydrogel sample. This yielded nine slides of soft hydrogel sections and nine slides of stiffened hydrogel sections. Samples on slides were stored at -80 ºC until immunofluorescent staining.

Cryosections on glass microscope slides were allowed to dry completely before immunofluorescent staining. Sections were fixed with ice-cold acetone for 15 minutes, then rinsed with room temperature water to remove OCT. Samples were allowed to dry completely then outlined with a hydrophobic pen. Sections were permeabilized with 0.2% Triton X-100 (Fisher Bioreagents) in PBS (Hyclone Laboratories, Inc., Logan, UT) for 10 minutes, then blocked with 5% bovine serum albumin (BSA; Fisher Scientific) in PBS for one hour at room temperature. Sections were incubated overnight at 4 ºC in primary mouse anti-human αSMA monoclonal antibody (Fisher Scientific catalog no.

MA5-11547) diluted 1:250 in immunofluorescent (IF) solution containing 3% BSA and 0.1% Tween 20 (Sigma) in PBS. After overnight primary antibody incubation, sections were rinsed three times in IF solution and incubated with secondary goat anti-mouse Alexa Fluor 555 antibody (Fisher Scientific) diluted 1:250 in IF solution with two drops of ActinGreen 488 ReadyProbes (Fisher Scientific) per mL of secondary antibody solution for one hour at room temperature protected from light. Sections were rinsed three times in IF solution and incubated with 300 nM DAPI (BioLegend, San Diego, CA) in deionized water for 15 minutes at room temperature protected from light. Sections were rinsed three times in deionized water protected from light, then cover-slipped and mounted with Prolong Gold Antifade Reagent (Fisher Scientific). Mounted slides were stored protected from light at -80 ºC until imaging.

Cryosectioned and immunofluorescent-stained hydrogels were imaged on an Olympus BX63 upright microscope. Three random cryosections per slide were imaged with DAPI, FITC, and TRITC filters under a 10× objective. Images were analyzed with ImageJ software (NIH). Fibroblast activation was quantified as the percentage of αSMA-positive cells.

### Assessment of Proliferation by Immunofluorescent Staining

Constructs were 3D-bioprinted with HPAAFs as described above, placed in the incubator overnight, and LifeSupport gelatin microparticle support bath was exchanged for 1% FBS activation media the following day (day 1). Media was changed on day 4. On day 6, 2.2 mM LAP was added to half of the samples and 10 µM 5-ethynyl-2’-deoxyuridine (EdU) solution from a Click-iT Plus EdU Cell Proliferation Kit (cat. no. C10637; Invitrogen, Waltham, MA) was added to the other half according to manufacturer instructions. All samples were incubated overnight. On day 7, samples incubated with LAP were stiffened as described above and samples incubated with EdU were fixed according to manufacturer instructions. On day 8, stiffened samples were incubated with 10 µM EdU solution overnight. Stiffened samples incubated with EdU were fixed according to manufacturer instructions on day 9.

Soft and stiffened 3D-bioprinted constructs containing EdU were processed for imaging according to manufacturer instructions. Images were acquired on a Zeiss LSM780 confocal microscope. Maximum intensity projections were created from 100-µm z-stacks acquired at three random fields of view per sample. HPAAF proliferation was measured by counting cells positive for EdU and normalizing to total cell number determined by Hoechst nuclear counterstain.

### Statistical Methods

Statistical analyses of sample means for two groups were performed with Mann-Whitney U tests. Statistical analyses of sample means for three or more groups were performed with one-way ANOVA tests followed by Tukey’s honest statistical difference (HSD) test for multiple comparisons. 3D construct viability results were compared with two-way ANOVA and Tukey’s HSD test to determine statistical significance. All statistical analyses were conducted in GraphPad Prism software (GraphPad Software, Inc., San Diego, CA) Hydrogel formulations were determined by a custom DOE generated with JMP software (SAS, Cary, NC). The resulting modulus and metabolic activity data were entered back into the custom DOE and analyzed with a least-squares regression model to determine the most significant factors and plot the predicted response.

## Results

### Hydrogel Formula Testing

A DOE approach identified seven different hydrogel formulations for evaluation of mechanical properties and cell metabolic activity. Each formulation contained an eight-armed PEG-αMA macromer with a molecular weight (MW) of 10 kg mol^−1^, two crosslinkers: a short dithiol crosslinker, DTT (MW = 154.25 g mol^−1^) and an MMP2-degradable peptide crosslinker (KCGGPQGIWGQGCK) to enable fibroblast remodeling, and a peptide sequence that mimics the adhesion protein fibronectin (CGRGDS; 2mM) to facilitate cell attachment.^[43]^ Reacting this system off-stoichiometry (0.375 [thiol]:[ene]) enabled fabrication of an initially soft hydrogel and sequential, light-initiated stiffening in the presence of lithium phenyl-2,4,6-trimethylbenzoylphosphinate (LAP) photoinitiator.^[35]^ Soft hydrogels exhibited elastic modulus (E) values ranging from 0.72 ± 0.12 kPa to 8.50 ± 0.53 kPa, while the elastic moduli of stiffened samples ranged from 4.80 ± 0.53 kPa to 20.55 ± 1.26 kPa (**Fig 1A**). The four formulations with soft modulus values within the range of healthy human pulmonary vascular ECM (1-5 kPa) and stiffened moduli that replicated the mechanical properties of hypertensive human vascular ECM (E = 10-30 kPa)^[8]^ were selected for additional 2D testing of cellular responses to the biomaterials prior to 3D bioprinting for viability and activation experiments. These four formulations were seeded with HLFs in 2D, and metabolic activity (analogous to proliferation)^[44-46]^ was monitored over time using a PrestoBlue Cell Viability assay. Results demonstrated increasing levels of cellular proliferation over all seven days of culture, and up to a seven-fold increase in metabolic activity at day 7 (**Fig 1B**). The best hydrogel formulation was defined as the platform that most accurately reproduced the healthy and pathological mechanical properties of pulmonary vascular ECM while maximizing fibroblast metabolic activity.

**Figure 1.**
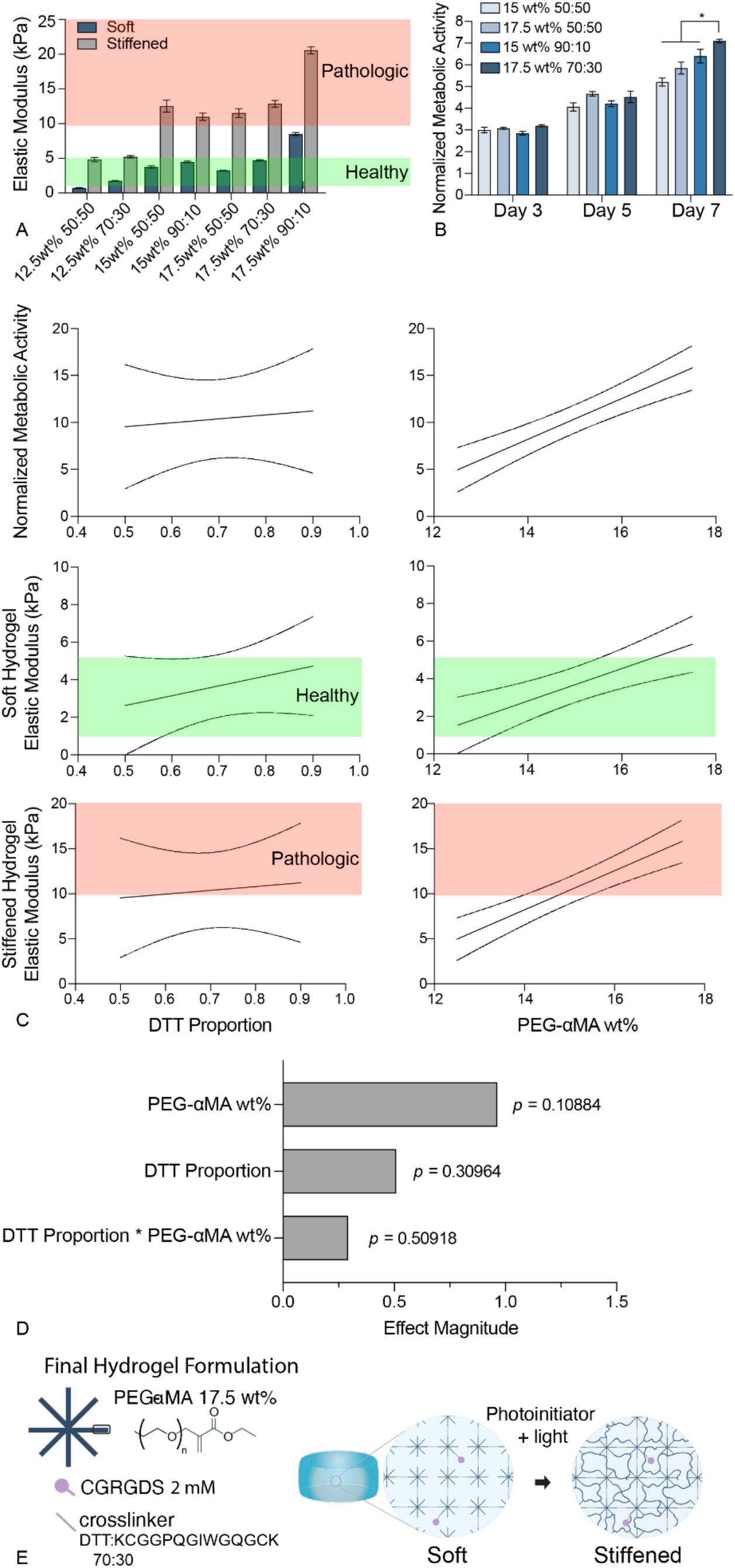
PEG-αMA hydrogel formulation selection. **A)** Rheological measurements of hydrogels with various compositions showed that several formulations have elastic moduli (E) that fall within the range of healthy physiologic pulmonary vasculature (1-5 kPa) when soft and rise to pathologic ranges (>10 kPa) when stiffened. Hydrogel formulations are presented as weight percent (wt%) of PEG-αMA followed by ratio of DTT:MMP2-degradable crosslinker, all with a ratio of [thiol]:[ene] of 0.375 for initial soft hydrogel fabrication. Columns represent mean ± SEM, n = 3. **B)** Human lung fibroblast (HLF) metabolic activity measured by PrestoBlue Cell Viability assay on various hydrogel formulations out to seven days of culture. Cells were cultured on 2D hydrogels and metabolic activity was normalized to data from day 1. Columns represent mean ± SEM, n = 3. *, *p* < 0.05, ANOVA, Tukey HSD. **C)** Design of experiments (DOE) effects analysis of DTT proportion and PEG-αMA wt% of hydrogel formulation on cell metabolic activity, soft hydrogel elastic modulus, and stiffened hydrogel elastic modulus. Data presented as predicted values for each outcome with 95% confidence interval. **D)** Effect magnitudes of DOE inputs on best hydrogel formulation. The best hydrogel formulation was determined as that which maximizes cell metabolic activity, exhibits a soft elastic modulus of 1-5 kPa, and a stiffened elastic modulus >10 kPa. **E)** Final hydrogel formulation determined by DOE analysis and schematic of hydrogel stiffening mechanism.

Both DOE inputs, DTT proportion and PEG-αMA wt%, correlated positively with hydrogel elastic modulus and cell proliferation. PEG-αMA wt% strongly influenced the elastic modulus results – hydrogel conditions with low PEG-αMA wt% did not achieve elastic modulus values high enough to replicate pathologic human tissue elastic modulus, whereas hydrogels with high PEG-αMA wt% exceeded the elastic modulus of healthy human tissue (**Fig 1C**). Effect analysis of DOE inputs (PEG-αMA wt% and DTT proportion) showed that PEG-αMA wt% was the most significant factor in these statistical metrics (*p* = 0.11, least-squares regression model; **Fig 1D**).

Hydrogel formulation desirability maximization through least-squares regression analysis of a custom DOE in JMP determined that a 17.5 wt% PEG-αMA hydrogel with a 70:30 ratio of DTT:MMP2-degradable peptide crosslinker and a [thiol]:[ene] ratio of 0.375 could make initially soft hydrogels (E = 4.7 ± 0.09 kPa) that could be stiffened to mimic pathogenic tissue (E = 12.8 ± 0.47 kPa), all while retaining high levels of cell proliferation (seven times greater at day 7 than day 1), which is significantly greater (*p* < 0.05, one-way ANOVA and Tukey HSD) than other formulations with similar elastic moduli (**Fig 1E**). This hydrogel formulation was used in all subsequent experiments.

### Dynamic Control of Hydrogel Mechanical Properties

The PEG-αMA hydrogel formulation that best achieved healthy and pathologic E values while supporting cell metabolic activity, identified through DOE analysis, was further evaluated for spatiotemporal control over mechanical properties. *In situ* rheology was conducted during both steps of the dual-polymerization process. Shear modulus measurements were collected beginning with liquid polymer precursor, during the Michael addition reaction, and throughout photoinitiated hydrogel stiffening. Base-catalyzed Michael addition resulted in an initial soft polymer (E = 3.82 kPa).

Photoinitiated homopolymerization caused hydrogels to stiffen to E = 73.18 kPa (**Fig 2A**) in the unswollen state. PEG-αMA hydrogels also demonstrated hydrolytic stability compared to PEG-MA hydrogels. Traditional PEG-MA hydrogels contain an ester linkage between the polymer backbone and the MA functional group. This is a preferential site for hydrolysis and leads to hydrogel degradation during long-term cell culture. PEG-αMA connects MA to the PEG backbone on the opposite side of the carbonyl allowing hydrolysis to occur without affecting the crosslinked polymer network.^[27]^ Stiffened PEG-MA and PEG-αMA hydrogel samples were soaked in PBS at 37 °C to simulate cell culture conditions and the elastic modulus was measured at days 1, 10, 20, 40, and 60. Initial Day 1 elastic modulus values did not differ between PEG-αMA and PEG-MA hydrogels (*p* > 0.05, nonparametric t-test). However, the elastic modulus measurements for PEG-MA hydrogels dropped quickly over time, and those samples showed significantly lower elastic modulus compared to PEG-αMA hydrogels at days 20, 40, and 60 (*p* < 0.05, nonparametric t-test). These results showed that PEG-αMA was not as susceptible to hydrolytic cleaving of crosslinks as PEG-MA and reinforced the use of PEG-αMA for long-term cell culture experiments performed to study chronic fibrotic diseases like PAH (**Fig 2B**).

**Figure 2.**
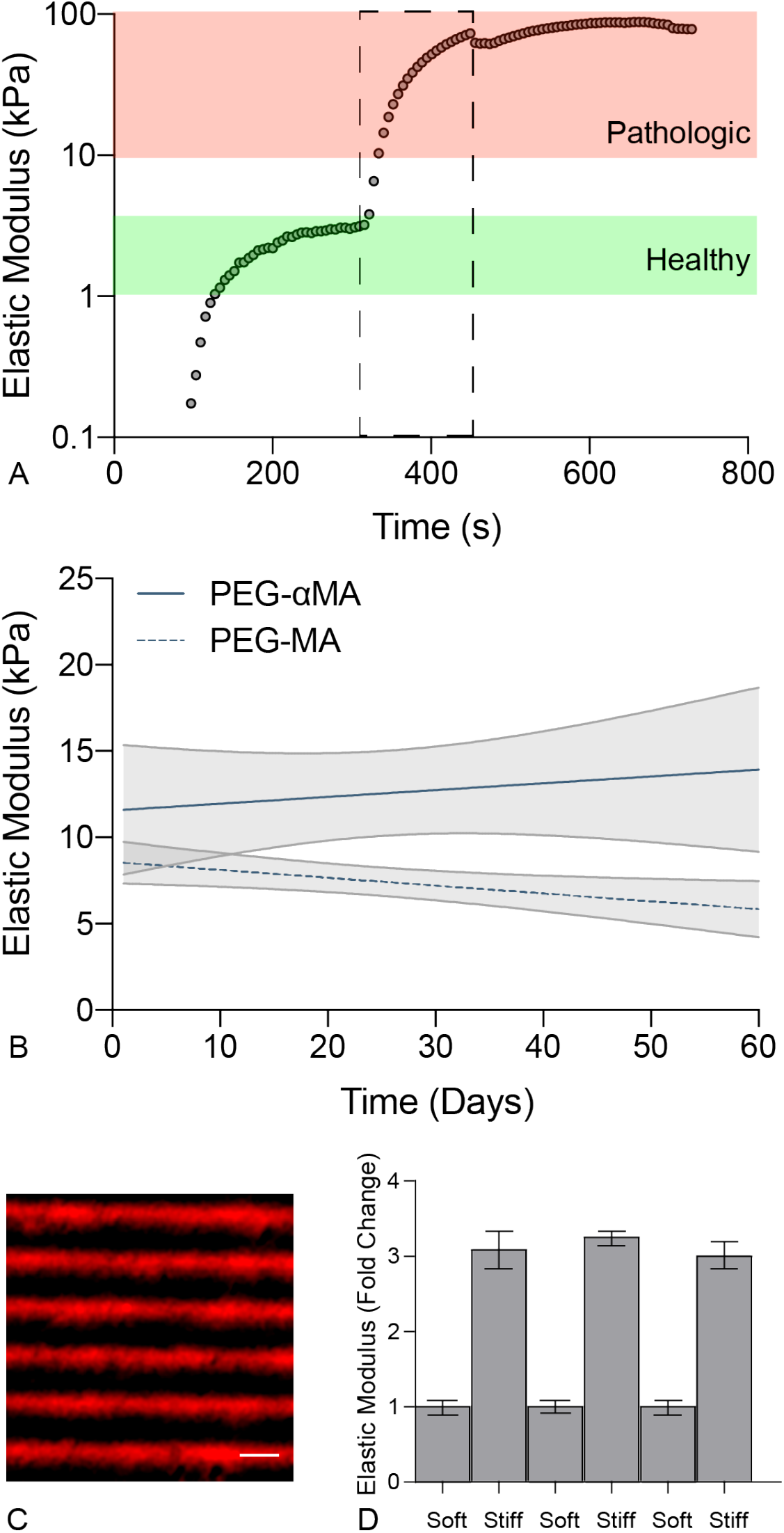
Spatial and temporal control over PEG-αMA hydrogel mechanical properties. **A)** Real-time rheological measurements of PEG-αMA elastic modulus during polymerization. The hydrogel underwent base-catalyzed initial polymerization to reach healthy pulmonary artery mechanical properties. Dotted box represents ultraviolet (UV) light-initiated homopolymerization, during which the hydrogel stiffens and achieves elastic modulus values seen in pulmonary hypertension pathology. **B)** Hydrolytic stability of stiffened PEG-αMA hydrogels compared to PEG-MA hydrogels. Lines represent mean elastic modulus, shaded regions represent 95% confidence interval, n = 4. **C)** Fluorescent microscopy image of a 2D PEG-αMA hydrogel with fluorescently labeled stiffened regions (red). Scale bar = 100 µm. **D)** Mechanical properties of 100-µm stiffened stripes on 2D PEG-αMA hydrogels as measured by atomic force microscopy. Bars represent mean elastic modulus ± SEM from two hydrogel samples with three stripes measured per sample and 15 measurements per stripe.

Next, spatial control over PEG-αMA hydrogel stiffening was demonstrated by exposing hydrogels to ultraviolet (UV light through a micropatterned photomask. Hydrogels were soaked in a fluorescent dye with the capacity to bind to αMA end groups during the photopolymerized stiffening reaction. UV light was delivered through a chrome-on-quartz photomask with 100-µm transparent stripes (quartz) separated by 100-µm opaque stripes (chrome), and the dye conjugated with the hydrogel wherever stiffening occurred. Fluorescent microscopy revealed discrete stiffened regions on the surface of a 2D hydrogel, represented by stiffened red stripes separated by soft non-fluorescent stripes (**Fig 2C**). Atomic force microscopy (AFM) measurements of photopatterned 2D hydrogel samples demonstrated distinct differences in elastic modulus between 100-µm soft stripes and 100-µm stiffened stripes. The average elastic modulus of stiffened stripes was 3.12 ± 0.07 times greater than that of soft areas (**Fig 2D**). These results demonstrated that PEG-αMA hydrogel stiffness was temporally patterned with a high degree of spatial control in 2D.

### Bioink Characterization

FRESH is a 3D bioprinting technique that allows precise creation of soft hydrogel constructs. PEG-αMA polymerizes through a base-catalyzed Michael addition reaction – the bioink remained aqueous in an acidic environment and become solid in a basic environment.^[27]^ Therefore, pH of the bioink was acidic (pH = 6.2) during extrusion to prevent gelation in the syringe. Conversely, the pH of the gelatin microparticle support bath remained basic (pH = 9.0) to initiate polymerization (**Fig S1A**). Cell metabolic activity was assessed at the acidic pH of the bioink and the basic pH of the gelatin microparticle support bath to ensure that fibroblasts would survive the printing and polymerization processes. PrestoBlue Cell Viability assays demonstrated no significant differences in metabolic activity of cells exposed to media at pH 6.2, pH 7.0, and pH 9.0 for 30 minutes (**Fig S1B**). These results showed that fibroblasts were as proliferative at the altered pH values used in the bioprinting process as at a normal culture pH of 6.9-7.3 over the period required for 3D bioprinting and polymerization (*p* > 0.05 by one-way ANOVA and Tukey HSD). To ensure printability and uniform cellular encapsulation, bioinks exhibited two key properties: 1) shear-thinning behavior, and 2) an initial viscosity value that prevents cell settling within the syringe on the 3D bioprinter.^[39, 47]^

The addition of small amounts of nonreactive PEO to the dynamic biomaterial produced shear-thinning rheological properties, i.e., a viscosity of 13.74 Pa at a shear rate of 0.1 s^−1^, descending to a viscosity of 1.07 Pa at a shear rate of 1,000 s^−1^. By contrast, unaltered hydrogel precursor had a viscosity of 0.004 Pa at a shear rate of 0.1 s^−1^, a value which remained constant as shear rate increased, with hydrogel precursor reaching a viscosity of 0.007 Pa at 1,000 s^−1^ shear rate (**Fig S1C**). A cell settling assay was performed to assess the capacity of the PEG-αMA bioink to retain an even distribution of cells throughout the syringe during the 3D bioprinting process. HLFs were labeled with CellTracker Green CMFDA fluorescent dye and suspended in PEG-αMA prepolymer solution with or without 2.5 wt% PEO (termed bioink or hydrogel, respectively). These cell solutions were loaded into cuvettes and placed vertically for 90 minutes to simulate the orientation of cells in the bioprinting syringe. Microscopic images of cell solutions in cuvettes were divided vertically into quadrants and the number of cells in each quadrant was counted.^[47]^ The bioink containing PEO showed a significantly lower number of cells in quadrant 1, the bottom of the cuvette, with 22 ± 2% of cells at the bottom of the cuvette in the bioink versus 31 ± 2% of cells at the bottom of the cuvette in unaltered hydrogel precursor solution, which are significantly different values (*p* < 0.05 by one-way ANOVA and Tukey HSD; **Fig S1D-E**). PEO-containing PEG-αMA bioink was shear-thinning, sustained an evenly distributed population of cells during 3D bioprinting and was used in all subsequent 3D bioprinting experiments.

### Bioprinted Construct Design

Confocal images of fluorescently labeled 3D-bioprinted hydrogels (**Fig 3A**) demonstrated that the FRESH 3D-bioprinting process produced cohesive hydrogel constructs for vascular modeling. These cylindrical structures contained pores induced by polymerization in the FRESH gelatin microparticle support bath. Such interstices in the polymer structure both mimic the microscale structure of the pulmonary arterial adventitia and provided a microenvironment amenable to cell infiltration and remodeling,^[48]^ especially in the presence of the MMP2-degradable peptide crosslinker employed in this hydrogel formulation. HPAAF viability in 3D pulmonary arterial adventitia models was measured by Live/Dead assays at days 1, 3, 7, 14, and 21 for each of four conditions combining two different wall thicknesses and two different HPAAF seeding densities (Table 3). Models with a 300-µm wall thickness and 4 × 10^6^ cells mL^−1^ showed a significantly greater percentage of live cells compared to all other conditions at day 7 and day 14 (*p* < 0.05, ANOVA and Tukey HSD; **Fig 3B**). This condition yielded 91 ± 2% live cells at day 7 and 85 ± 3% live cells at day 14 and was used for future experiments. Models with a 500-µm wall thickness and 4 × 10^6^ cells mL^−1^ had a significantly greater percentage of live cells at day 7 compared to tubes with 500-µm wall thickness and 4 × 10^5^ cells mL^−1^ (73 ± 4% versus 47 ± 5%, respectively; *p* < 0.05, ANOVA and Tukey HSD). Interestingly, the 500-µm wall thickness and 4 × 10^6^ cells mL^−1^ condition had a significantly lower percentage of live cells at day 21 compared to the most successful condition, tubes with 300-µm wall thickness and 4 × 10^6^ cells mL^−1^ (17 ± 7% vs. 56 ± 9%, respectively; *p* < 0.05, ANOVA and Tukey HSD). Percentage of live cells in 3D-bioprinted samples peaked at day 7 for all conditions, so this time point was used in future experiments (**Fig 3C**).

**Figure 3.**
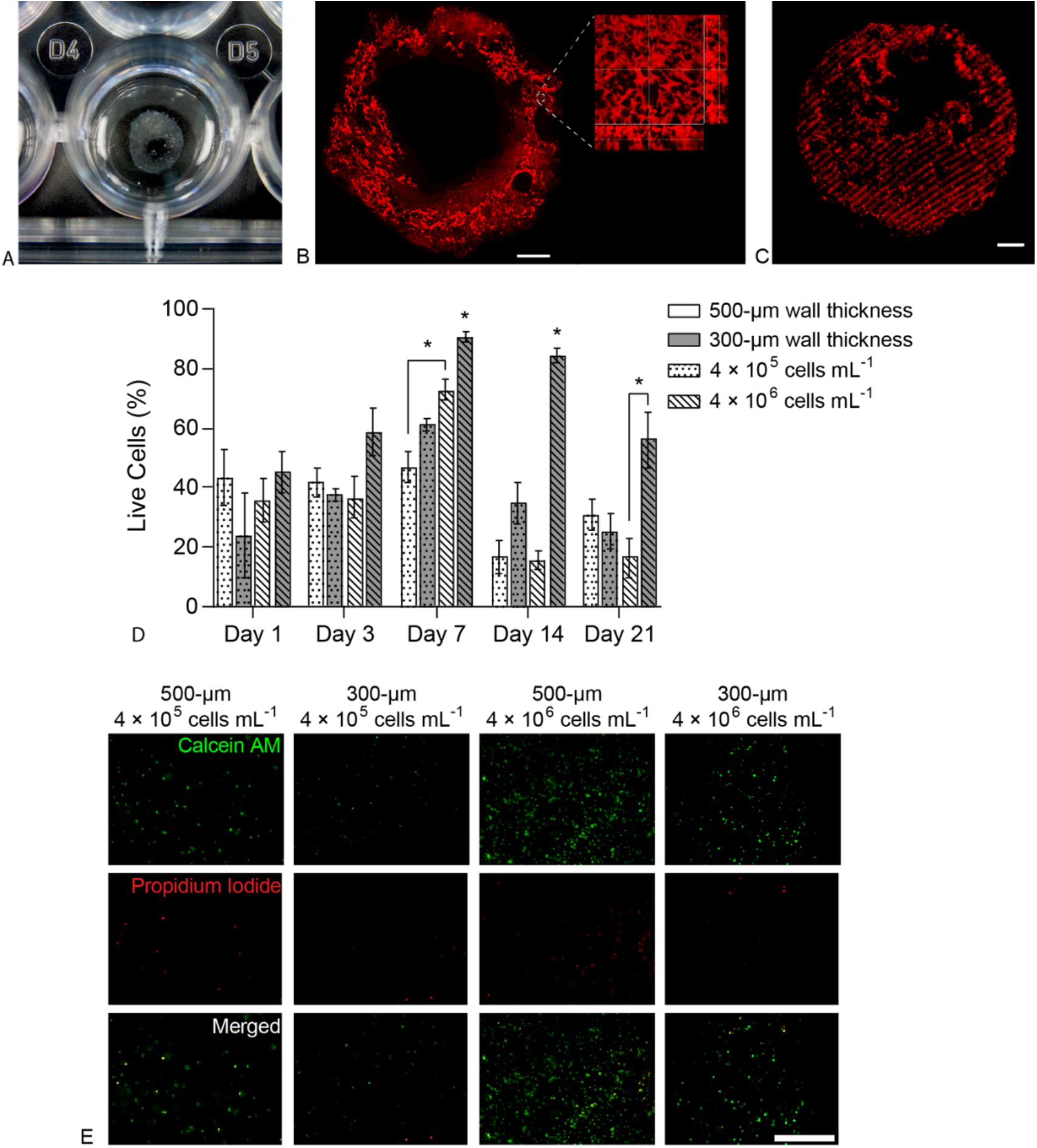
3D-bioprinted hydrogel structures support cell viability over time. **A)** Photograph of 3D-printed hydrogel structure in a 24-well plate. **B)** Maximum intensity projection of fluorescently labeled PEG-αMA 3D-printed hydrogel. Scale bar = 1mm. Higher magnification microscopy showed pores within the hydrogel structure induced by gelatin microparticles in the FRESH bioprinting support bath. **C)** 3D-printed PEG-αMA tube with fluorescently labeled stiffened regions imaged on a confocal microscope (100-µm z-stack displayed as a maximum intensity projection) showed spatial control over stiffening in 3D. Scale bar = 500 µm. **D)** HPAAF viability in 3D-bioprinted constructs measured by Live/Dead assays. Constructs with 300-µm thickness and 4 × 10^6^ cells mL^−1^ outperformed all other conditions at every time point. Viability peaked at day 7. This condition and time point were selected for future experiments. Columns show mean ± SEM, n = 3. *, *p* < 0.05, ANOVA, Tukey HSD. **E)** Representative confocal images of cells in 3D constructs stained with Live/Dead reagent at day 7, the time point with greatest overall viability. Calcein AM marked live cells in green, and propidium iodide marked dead cells in red. The right-most column shows that the best-performing condition had uniform cell distribution and a high percentage of live cells. Scale bar = 500 µm.

### Fibroblast Activation

The influence of dynamic changes in microenvironmental stiffness on fibroblast activation was assessed by measuring proliferation, αSMA expression, and extracellular matrix remodeling. Primary male HPAAFs were cultured in 3D-bioprinted PEG-αMA hydrogels, half of the samples were stiffened on day 7, and activation was compared between soft and stiffened samples. HPAAFs cultured in stiffened 3D hydrogels showed significantly greater expression of αSMA, a marker of fibrotic activation, compared to HPAAFs cultured in soft 3D hydrogels. Stiffened hydrogels produced 88 ± 2% αSMA-positive cells, compared to 65 ± 4% in soft hydrogels (*p* < 0.05 by Mann-Whitney U test; **Fig 4A**). This difference in αSMA-positive cells could be clearly seen in images of hydrogel cryosections (**Fig 4B**). Our lab has verified that the increased expression of αSMA protein in stiffened hydrogels compared to soft hydrogels quantified through image analysis was consistent with αSMA protein expression measured by Western blot analysis (unpublished data). Additionally, HPAAFs cultured in stiffened 3D constructs showed a marked increase in proliferation over cells cultured in soft hydrogels as measured by EdU-positive percentage. HPAAFs in stiffened hydrogels were 66 ± 6% positive for EdU, a marker of proliferation, compared to 39 ± 6% of cells in soft hydrogels (*p* < 0.05 by Mann-Whitney U test; **Fig 4C**). HPAAFs that synthesized new DNA during the overnight EdU incubation can be seen in the immunofluorescent images of stained antibodies in soft and stiffened 3D hydrogel constructs (**Fig 4D**).

**Figure 4.**
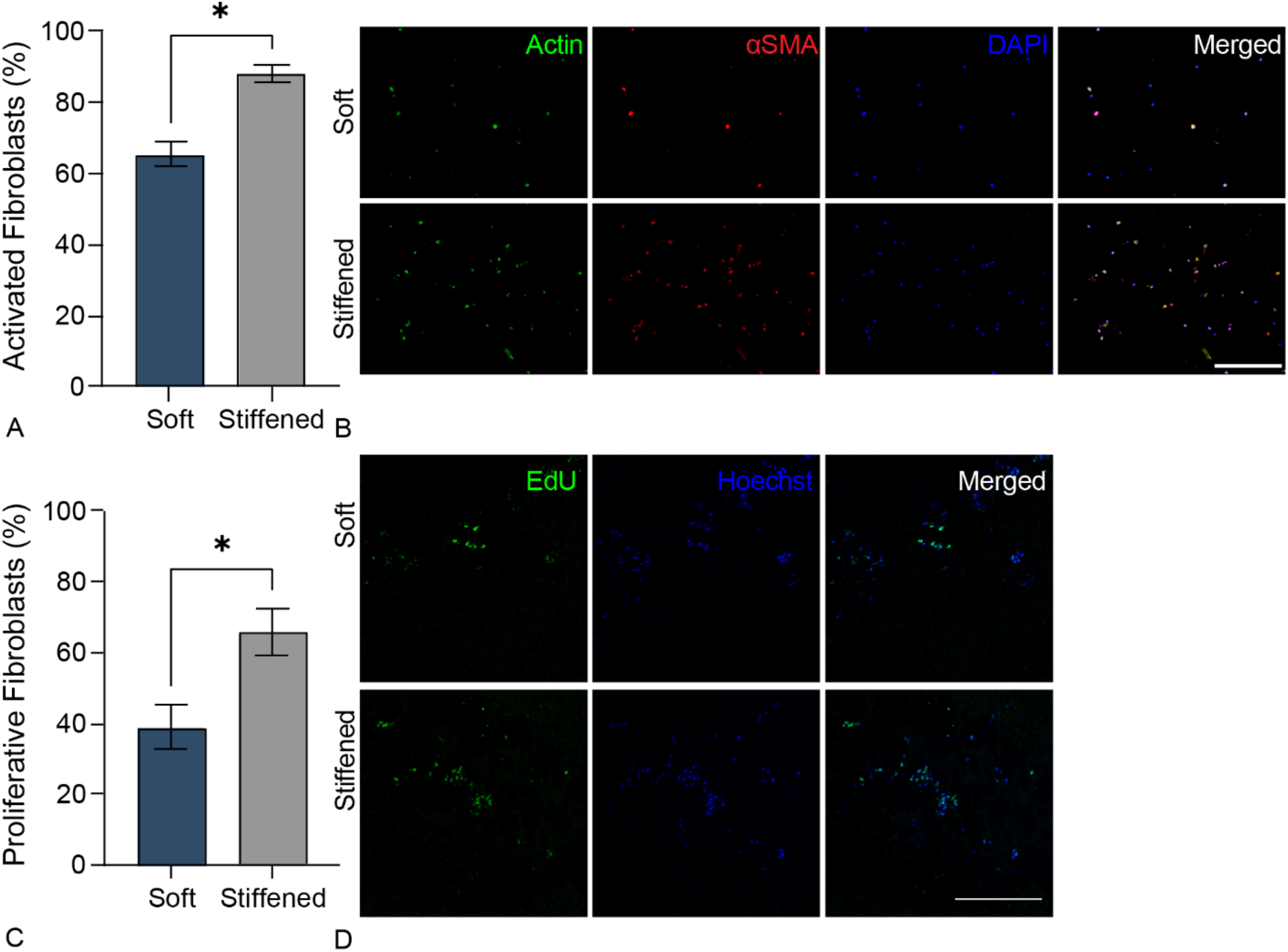
Fibroblast activation in 3D-bioprinted models of pulmonary arterial adventitia. **A)** Fibrotic activation in soft and stiffened 3D hydrogels measured by αSMA expression. HPAAFs in stiffened constructs were significantly more positive for αSMA than cells in soft constructs. Columns represent mean ± SEM, n = 3. *, *p* < 0.05, Mann-Whitney U test. **B)** Representative confocal images of immunostaining for αSMA, actin, and DAPI in soft and stiffened 3D hydrogels. HPAAFs in stiffened constructs showed more prevalent αSMA immunofluorescence than cells in soft constructs. Scale bar = 250 µm. **C)** Fibroblast proliferation in soft and stiffened 3D bioprinted constructs measured by EdU positivity. HPAAFs in stiffened constructs were significantly more positive for EdU than cells in soft constructs. Columns represent mean ± SEM, n = 3. *, *p* < 0.05, Mann-Whitney U test. **D)** Representative confocal images of immunostaining for EdU and Hoechst dye in soft and stiffened 3D hydrogels. HPAAFs in stiffened constructs showed more prevalent EdU immunofluorescence than cells in soft constructs. Scale bar = 300 µm.

HPAAFs in 3D hydrogel substrates remodel their extracellular microenvironment, as shown by rheology of cellularized hydrogels, MMP2 secretion, and collagen deposition in 3D-bioprinted constructs. Hydrogels containing embedded HPAAFs demonstrated increases in elastic modulus over time up to the day 7 photoinitiated homopolymerization stiffening, with 2-fold greater elastic modulus at day 7 compared to day 1 without stiffening, and 3-fold greater elastic modulus with homopolymerization compared to day 1. Soft samples at day 9 still showed twice the elastic modulus as day 1 samples, but stiffened day 9 samples had elastic moduli only 1.5 times greater than day 1, showing a decrease in elastic modulus between day 7 and day 9 (**Fig S2A**).

Enzyme-linked immunosorbent assays (ELISAs) of 3D-bioprinted constructs showed that HPAAFs consistently secreted MMP2. Supernatant from samples at day 1 contained 19.0 ± 1.0 ng mL^−1^ MMP2, which increased to 26.0 ± 1.5 ng mL^−1^ at day 3 and peaked at 24.5 ± 1.0 ng mL^−1^ at day 7 before stiffening. Samples at day 9 secreted lesser quantities of MMP2, with supernatant from soft samples containing 17.4 ± 0.7 ng mL^−1^ and supernatant from stiffened samples containing 17.8 ± 0.8 ng mL^−1^ (**Fig S2B**). Cells also deposited collage, as visualized by Picro Sirius Red stain. This collagen was in the pericellular space and was present from day 1 all the way through the end of culture in soft and stiffened 3D-bioprinted constructs (**Fig S2C**).

## Discussion

The experiments performed here combined a phototunable biomaterial with a 3D-bioprinting technique to evaluate pulmonary artery adventitial fibroblast activation in a dynamic 3D microenvironment. The PEG-αMA formulation was designed so that initial hydrogel crosslinking resulted in a platform with elastic modulus representative of the non-pathologic vascular environment, and the secondary crosslinking caused stiffening to create a hydrogel representative of the pathologic adventitia elastic modulus in PAH (1-5 kPa and 10-30 kPa, respectively).^[8]^ A DOE approach maximized desirability of the hydrogel formula to create a 3D microenvironment with physiological mechanical properties not commonly achieved by traditional cell culture techniques or 2D organ-on-a-chip designs that use stiffer substrates.^[49]^ Additionally, this 3D hydrogel presented cells with a more physiologically accurate microenvironment than similarly formulated 2D hydrogels – the 3D culture system allowed adhesion on all surfaces of the cells, encouraged spreading and migration in all three dimensions, and did not force an apical-basal polarity.^[50, 51]^ The PEG-αMA material used here allowed precise spatial control over the stiffening reaction as measured by AFM (Fig 2E). The biomaterials field has widely employed UV light to induce microenvironmental stiffening or softening to study fibroblast activation.^[18, 22, 52]^ Ruskowitz and DeForest analyzed fibroblast response to exposure to 365-nm light – the same wavelength used in this study – and found unchanged proliferation rates, no induction of apoptosis, and unchanged proteome.^[53]^ This spatial control, combined with temporal control of stiffness via initiation of the photopolymerization reaction, created a versatile and reproducible platform to study the role of fibroblast activation in PAH.

The combination of thiol-ene chemistry and pH-driven polymerization produced a novel system to study cells in 3D conditions. Cell populations in hydrogel structures remained proliferative through pH-initiated polymerization over an extended print time (Fig S1B). This reinforces prior work showing that changes in the microenvironmental pH of 3D-bioprinted constructs do not severely hamper proliferation of cell constructs.^[54]^ A recent study showed similar cell survival in a FRESH system that polymerized collagen through precise control of solution pH.^[48]^ The PEG-αMA system produces a unique FRESH 3D-bioprinted culture platform by taking advantage of different chemistries and crosslinking reactions.

This fibrotic model builds on research regarding lung-mimetic bioprinted architecture^[55]^ and provides a hydrogel platform that can be used to study the influence of 3D culture conditions on fibroblast behavior. The 3D-bioprinted hydrogel microstructure contains interstices reminiscent of decellularized human vascular adventitia.^[56]^ The two-step polymerization method implemented here enabled evaluation of cellular proliferation and phenotype in 3D-bioprinted constructs before and after stiffening. Viability was maximized by tailoring construct wall thickness and cell density through a full-factorial approach. Constructs had wall thicknesses of 300 or 500 µm, replicating dimensions reported by Dai et al. in their measurements of third to fourth generation human pulmonary arteries and Li et al. in their research regarding 3D-bioprinted vessels.^[40, 41]^ Likewise, the bioink was seeded with fibroblasts 4 × 10^5^ or 4 × 10^6^ cells mL^−1^, densities established by Li et al. and Zhang et al. in recent 3D vascular cell studies.^[41, 42]^ Live/Dead assays showed that constructs with 300-µm wall thickness and 4 × 10^6^ cells mL^−1^ had the greatest viability out to 21 weeks, with a peak in percentage of live cells at day 7 (Fig 3B-C). The increased cell viability in thin-walled bioprinted constructs is likely due to improved transport of media and metabolites through the hydrogel compared to thicker walls. Poor viability at day 1 could be caused by the extrusion pressure of the bioprinting syringe head, which must be great enough to deposit the shear-thinning bioink but may cause cell death. These results established the most desirable 3D-bioprinted construct parameters and experimental time points for fibroblast activation studies.

The argument that adventitial ECM stiffening may have a driving role in PAH progression is supported by the identification of mechanosensitive pathways that drive disease progression *in vivo*, such as the YAP/TAZ-miR-130/301 circuit.^[7]^ Furthermore, activated adventitial fibroblasts are implicated in induction of a proinflammatory and profibrotic macrophage phenotype in PAH.^[57]^ For this reason, *in vitro* models have been valuable for probing the connection between ECM stiffness and PAAF activation.

Previous studies have aimed to investigate this connection. Wang et al. cultured rat PAAFs on polyacrylamide hydrogels of varying stiffness; smooth muscle actin (SMA) expression measured by corrected total cell fluorescence (CTCF) was not significantly different in these cells when grown on 3 kPa or 10 kPa substrates.^[58]^ The data reported by that research team contrast the results of this study, which found a significant difference in human PAAF expression of αSMA when cultured on hydrogels with similar differences in stiffness. However, a few key factors differentiate these two studies. The bioprinted model described in this study used human cells and incorporated an additional dimension, encapsulating human PAAFs in 3D hydrogels instead of seeding rat PAAFs on top of 2D hydrogels. Also, protein expression was measured in cells contained in a dynamically stiffened substrate that had previously grown in a soft microenvironment, instead of comparing two cell populations grown on static soft or stiff hydrogels.

This report describes a new biomaterial for 3D-bioprinting dynamic cell culture platforms to study the effects of 3D culture and matrix modulus on human adventitial fibroblast activation. This dynamic hydrogel incorporated an MMP2-degradable peptide to facilitate cellular remodeling of the microenvironment. Levels of various MMPs are generally increased in patients with PAH, and MMP2 has shown clinical significance in disease prognosis.^[59, 60]^ Adventitial fibroblasts produce MMP2 during development and during dysregulated remodeling of the vasculature, such as that occurring in PAH.^[32]^ Including an MMP2-degradable crosslinker allowed cell spreading prior to stiffening facilitating evaluation of the fibroblast-to-myofibroblast transition. Other synthetic 3D environments, which encapsulated cells in non-degradable hydrogels, restricted cell spreading and therefore showed significantly lower αSMA expression in human pulmonary fibroblasts cultured in stiff constructs compared to soft constructs.^[61]^ By contrast, results of the current study showed that stiffened hydrogels significantly increased expression of αSMA and EdU compared to soft hydrogels. The increase in HPAAF activation observed in stiffened 3D hydrogel constructs followed a week of culture, during which cells adhered and spread in a soft hydrogel substrate. This timeline differed from a previous study by Mabry et al. that showed decreased fibroblast expression of αSMA in 3D hydrogels following 48 hours of culture.^[62]^ Studies of HPAAF activation on 2D dynamically stiffening PEG-αMA hydrogels with biodegradable crosslinkers showed a similar increase in αSMA expression.^[63]^ The combination of MMP2-degradable crosslinks and long culture time produced a resilient 3D hydrogel microenvironment that allowed cell spreading and activation.

PAH is a progressive disease that ultimately leads to death – recently published longitudinal studies show that hypertensive proximal pulmonary arteries stiffen over the course of months or years.^[64, 65]^ A platform that allows observation of cell-ECM interactions in a pathologically stiff microenvironment after initially mimicking the early healthy mechanical properties is a valuable research tool. This 3D-bioprinted model exhibited a step change in elastic modulus (Fig 2A) that does not accurately represent the gradual increase in elastic modulus observed in clinical subjects. However, the near-instantaneous stiffening of these hydrogel platforms can accelerate experiments that would take much longer to conduct *in vivo* or through clinical data. Another limitation of this study is that it only included fibroblasts. This complements a recent study that constructed a 3D vascular graft with endothelial cells and vascular smooth muscle cells,^[66]^ but future studies could incorporate all three cell types pertinent to pulmonary vascular disease. Of course, long-term cell culture experiments require a hydrogel platform that will not break down in aqueous conditions. One study hypothesized that some state-of-the-art biomaterial pulmonary vascular mimics will soften over time.^[67]^ To mitigate this potential limitation, the tunable polymer network described in the current study employed hydrolytically stable αMA homopolymerized crosslinks that maintained stiffness in the fibrotic range out to 60 days (Fig 2B).

Future models of pulmonary vasculature with user-controlled, physiologic mechanical properties and patient-derived cells will enable researchers to probe the sequence of events that drives PAH. Follow-up research could build on this model of fibrotic activation, which accounted for the stiffening observed in hypertensive pulmonary vasculature and added additional factors that may contribute to PAH pathogenesis.^[12]^ A layer of biomaterial containing pulmonary artery smooth muscle cells could be 3D-bioprinted inside a layer of biomaterial with HPAAFs, and pulmonary artery endothelial cells could be flow-seeded onto the inside of this multicellular construct. This procedure would create a vascular model with cell layers appropriate to the tunica adventitia, tunica media, and tunica intima. This 3D combination of cellular layers would better replicate blood vessels than 2D hydrogel models. Flow conditions could also be incorporated to replicate the passage of blood through a healthy or diseased pulmonary artery. This would add another key dynamic aspect to the culture model described here that would represent human physiology better than 2D culture methods. Importantly, PAH affects female patients more frequently and at younger ages than male patients, although male patients exhibit more severe outcomes and decreased survival. These differences likely arise from the complex role of sex hormones, including the estrogen paradox, on pathogenesis. Experiments conducted in this study could be reproduced with female HPAAFs and sex-specific human serum to replicate *in vitro* the sex differences observed in PAH *in vivo*.^[68]^

## Conclusion

In summary, we have designed a cell culture platform that more accurately mimics the native and pathologic pulmonary arterial adventitial microenvironment. Hydrolytically stable, 3D-bioprintable PEG-αMA supported two-step polymerization with a high degree of spatiotemporal control which enabled *in vitro* user-controlled matrix stiffening as measured by rheology and AFM. A DOE approach was used to identify a hydrogel formulation that produces initially soft hydrogels representative of healthy pulmonary adventitia that can be stiffened to mimic pathogenic tissue. These biomaterials produced 3D-bioprinted constructs with 300-µm wall thickness that replicated small pulmonary blood vessels. These constructs were used to probe the mechano-related dynamic response of HPAAFs. Cells cultured on stiffened hydrogels showed a greater percentage of αSMA-positive cells and EdU-positive cells, correlating to increased fibrotic activation. The results of this study give a close look at the effects of stiffening 3D microenvironments on fibrotic activation, providing researchers with a versatile new platform to study PAH and other fibrotic diseases of the pulmonary vasculature. The advanced biomaterial platform designed here will provide the foundation for blood vessel models of increasing complexity comprising multiple cell types that are cultured under flow to reveal novel mechanistic insights into reduction of human disease. The complete model provides a platform for testing and validating therapies for pulmonary hypertension, advancing translational research in this field.

## Declaration of Competing Interests

Chelsea Magin reports a relationship with Boulder iQ that includes consulting fees. Chelsea Magin has patent #PCT/US2019/012722 pending to University of Colorado. All other authors, Duncan Davis-Hall, Emily Thomas, and Brisa Peña, declare that they have no known competing financial interests or personal relationships that could have appeared to influence the work reported in this paper.

## Acknowledgements

We thank Dr. Adam Feinberg for hosting the 3D Bioprinting Open-Source Workshop at Carnegie Mellon University, where we learned the FRESH printing technique and built our 3D bioprinter. Thanks also to Dr. Steven Lammers for generously sharing his equipment while ours was in transit. Finally, thanks to Dr. Keith Neeves for his help with the F31 Predoctoral Award application that funded parts of this study, and for his feedback during the writing of this article.

This study was supported by the Ludeman Family Center for Women’s Health Research at the University of Colorado Anschutz Medical Campus to DDH and CMM; the Rose Community Foundation to DDH and CMM; the National Heart, Lung, and Blood Institute of the National Institutes of Health (NIH) under awards R01 HL080396 (CMM), R01 HL153096 (CMM), K25 HL148386 (BP), F31 HL151122 (DDH), and T32 HL072738 (DDH); the National Cancer Institute of the NIH under award R21 CA252172 (CMM); the National Science Foundation under award 1941401 (CMM); the Department of the Army under award W81XWH-20-1-0037 (CMM); and a Colorado Pulmonary Vascular Disease Research Award to DDH and CMM. This material is based upon work supported by the National Science Foundation Graduate Research Fellowship Program under Grant No. DGE 1841052 (ET).

## Data Availability

The raw and processed data required to reproduce these findings are available here: Magin, Chelsea; Davis-Hall, Duncan; Thomas, Emily; Peña, Brisa (2022), “3D-bioprinted, phototunable hydrogel models for studying adventitial fibroblast activation in pulmonary arterial hypertension”, Mendeley Data, V1, doi: 10.17632/vycw3v6c5m.1.

## Notes

https://doi.org/10.17632/vycw3v6c5m.1

