## Supplementary Materials for "3D-bioprinted, phototunable hydrogel models for studying adventitial fibroblast activation in pulmonary arterial hypertension"

### Supplementary Materials and Methods

#### *Viscosity Characterization*

For viscosity measurements, a hydrogel precursor solution containing 17.5 wt% PEG-αMA with a 0.375 molar ratio of 50:50 DTT:MMP-degradable crosslinkers relative to the PEG-αMA backbone was prepared in pH 8.0 HEPES with and without the addition of 2.5 wt% 400 kg $mol^{-1}$ PEO. Viscosity of both samples was measured with a 40-mm 1° cone with analysis in flow sweep mode from 0.1 to 1,000 1 $s^{-1}$ shear rate at 22.5 °C using a solvent trap on a DHR-2 rheometer.

#### *Cell Settling*

HLFs (passage 7) were labeled with CellTracker Green CMFDA Dye (Thermo Fisher Scientific, Waltham, MA). HLFs were suspended at 50,000 cells $mL^{-1}$ in 10 mM CellTracker Green CMFDA working solution prepared in serum free media (DMEM/F12 1:1 supplemented with 100 U $mL^{-1}$ penicillin, 100 μg $mL^{-1}$ streptomycin and 2.5 μg $mL^{-1}$ amphotericin B, Gibco, Waltham, MA). Labeled HLFs were incubated at 37 °C for one hour then CellTracker solution was removed with centrifugation at 300 g for five minutes, and cells were resuspended in media. Cell solution was suspended uniformly in the bioink (containing PEO) or hydrogel (without PEO) and loaded into a square spectrophotometer cuvette (Thomas Scientific Swedesboro, NJ). Cuvette stood upright for one hour then was laid flat and imaged with an Olympus BX63 fluorescent microscope.[39] Resulting images were divided into four vertical quadrants and the total cells per quadrant were counted using ImageJ.

*pH Metabolic Activity*

Glass coverslips were soaked in media overnight at 37 °C, then seeded with HLFs (passage 5) at 10,000 cells $cm^{-2}$ and incubated again at 37 °C overnight. PrestoBlue Cell Viability Reagent (Thermo Fisher Scientific, Waltham, MA) was added to each sample following manufacturer instructions, and fluorescence was read after 30 minutes of incubation at 37°C on a rocker to establish baseline metabolic activity. Media was removed from each sample and replaced with media at pH 6.2 or 9.0, prepared with the addition of HCl (Thermo Fisher Scientific, Waltham, MA) or NaOH (Thermo Fisher Scientific, Waltham, MA), or unadjusted media (n = 3). Samples were incubated at room temperature under these media conditions for 40 minutes. PrestoBlue Cell Viability Reagent was added to each sample, and fluorescence was read after 30 minutes of incubation at 37°C on a rocker.

*Rheological Characterization of Cellularized Hydrogels*

Hydrogel solution was prepared with 17.5 wt% PEG-αMA, a 0.375 molar ratio of 70:30 DTT:MMP2-degradable crosslinker, and 2 mM RGD pendant peptide in pH 8 HEPES. HPAAFs were embedded in this solution at a density of $4 \times 10^6$ cells $mL^{-1}$. Six samples were prepared according to the procedure under *Rheological Characterization* methods for analysis on days 1, 3, 7, and 9. An additional six samples were prepared for photoinitiated stiffening on day 7 and measurement on day 9. All samples were cultured in complete media as previously described in this manuscript. Elastic modulus was measured at days 1, 3, 7, and 9 as previously described in this manuscript. On day 6, 2.2 mM LAP was added to six samples of each condition for stiffening on day 7, with corresponding measurement of stiffened elastic modulus on day 9.

*Quantification of MMP2 Secretion by Enzyme-Linked Immunosorbent Assay (ELISA)*

Concentration of MMP2 secreted by HPAAFs in 3D-bioprinted constructs was measured using the Human MMP2 ELISA Kit (Abcam, Cambridge, UK). Samples were bioprinted as described, and supernatant was collected at days 1, 3, 7, and 9. Day 9 supernatant was collected from both soft samples and samples photostiffened on day 7 following incubation with 2.2 mM LAP. The ELISA was conducted according to manufacturer's instructions using supernatant diluted 1:2.

*Visualization of Collagen Deposition*

Collagen deposition by HPAAFs in 3D-bioprinted constructs was measured by Picro Sirius Red Stain Kit (Connective Tissue Stain; Abcam, Cambridge, UK). Sample were 3D bioprinted as described, and culture media was supplemented with 75 μg $mL^{-1}$ L-ascorbic acid (Sigma-Aldrich, St. Louis, MO) to encourage collagen maturation. Samples were harvested and cryosectioned at days 1, 3, 7, and 9. Day 9 samples included soft constructs as well as constructs photostiffened on day 7 after incubation with 2.2 mM LAP. Hydrogels were also created without HPAAFs to for comparison. Cryosectioned samples were stained with Picro Sirius Red Stain Kit per manufacturer's instructions, then rinsed with DI water and incubated with 300 nM DAPI for 15 minutes protected from light. Slides were rinsed again with DI water and mounted for imaging.

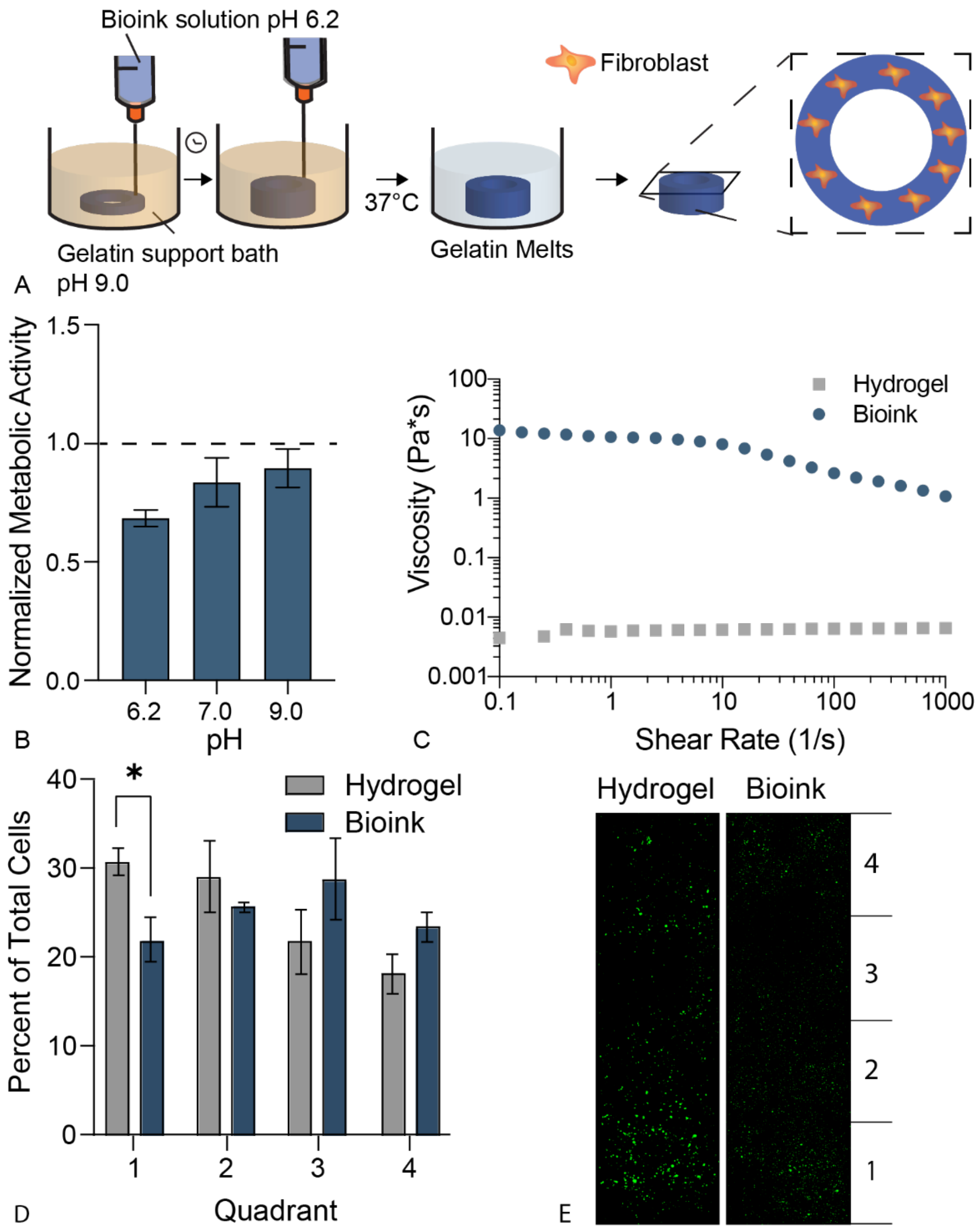


**Figure S1.** 3D bioprinting of PEG-αMA hydrogel bioinks. **A)** Schematic of FRESH 3D bioprinting. A slightly acidic bioink containing suspended cells was deposited by syringe into a slightly basic gelatin microparticle support bath using G-code generated from user-created 3D CAD models. The acidic bioink prevented polymerization during extrusion, and the basic support bath caused initial hydrogel polymerization. Physiologic temperature melted the gelatin support bath, leaving a cellularized 3D bioprint. **B)** HLF metabolic activity measured by PrestoBlue Cell Viability assays at various pH values used for the PEG-αMA bioink and gelatin microparticle support bath. **C)** Rheological measurements of material viscosity. Adding 2.5 wt% PEO to PEG-αMA hydrogel prepolymer provided shear-thinning properties that enabled stable suspension of cells in the syringe during printing while allowing extrusion through a narrow-gauge needle. **D)** Cell settling measured by suspending cells in hydrogel prepolymer or bioink containing PEO. Columns represent mean ± SEM, n = 3. *, $p < 0.05$, ANOVA, Tukey HSD. **E)** Image of HLFs in cell settling experiment labeled with CellTracker Green CMFDA. Quadrants shown at different cuvette depths were used to quantify results.

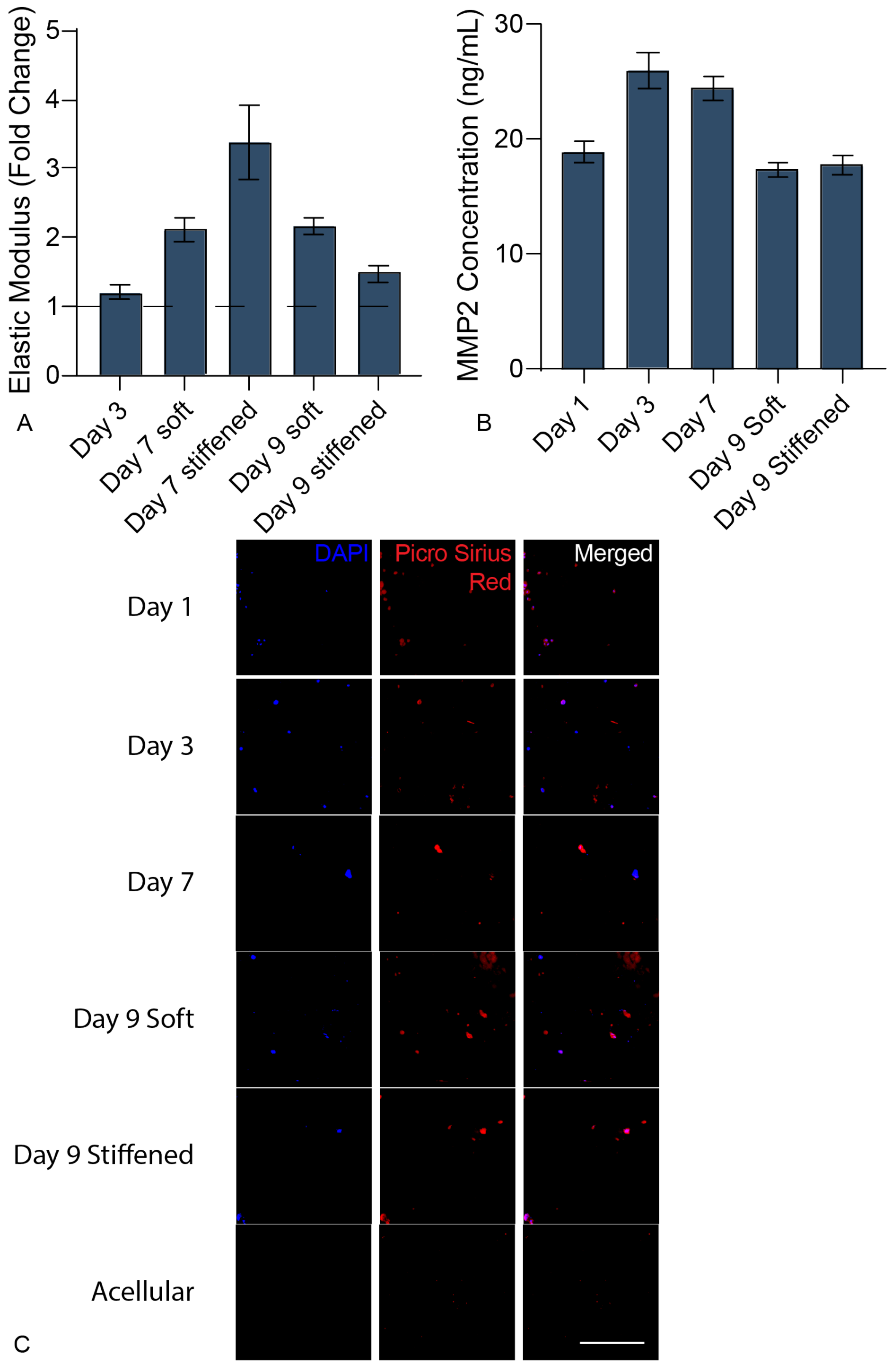


**Figure S2.** Characterization of matrix remodeling. **A)** Elastic modulus of cellularized 3D hydrogel samples normalized to day 1. Rheology showed increasing bulk elastic modulus over time, peaking at the day 7 photoinitiated homopolymerization stiffening event, with declining elastic modulus following. Columns represent mean ± SEM, n = 6. **B)** MMP2 concentration in supernatant of 3D-bioprinted constructs. HPAAFs secreted MMP2 continuously during culture, with decreased concentration after seven days in culture. Columns represent mean ± SEM, n = 3. **C)** Picro Sirius Red connective tissue stain shows pericellular deposition of collagen protein over the entire culture period. Scale bar = 200 μm.
